# α_2_-ADRENERGIC MODULATION OF I_h_ IN ADULT-BORN GRANULE CELLS IN THE OLFACTORY BULB

**DOI:** 10.1101/2022.09.27.509763

**Authors:** Ruilong Hu, Janam Shankar, Grant Z. Dong, Pablo S. Villar, Ricardo C. Araneda

## Abstract

In the olfactory bulb (OB), a large population of axon-less inhibitory interneurons, the granule cells (GCs), coordinate network activity and tune the output of principal neurons, the mitral and tufted cells (MCs), through dendrodendritic interactions. Furthermore, GCs undergo neurogenesis throughout life, providing a source of plasticity to the neural network of the OB. The function and integration of GCs in the OB is regulated by several afferent neuromodulatory signals, including noradrenaline (NA), a state-dependent neuromodulator that plays a crucial role in the regulation of cortical function and task-specific decision processes. However, the mechanisms by which NA regulates GC function are not fully understood. Here, we show that NA modulates hyperpolarization-activated currents (I_h_) via the activation of α_2_-adrenergic receptors in adult-born GCs (abGCs), thus directly acting on channels that play essential roles in regulating neuronal excitability and network oscillations in the brain. This modulation affects the dendrodendritic output of GCs leading to an enhancement of lateral inhibition onto the MCs. Furthermore, we show that NA modulates subthreshold resonance in GCs, which could affect the temporal integration of abGCs. Together, these results provide a novel mechanism by which a state-dependent neuromodulator acting on I_h_ can regulate GC function in the bulb.

## INTRODUCTION

The precise regulation of inhibitory circuits is an inherent component of sensory processing (Griffen & Maffei, 2014; Isaacson & Scanziani, 2011; Lledo et al., 2005). In the olfactory bulb (OB), the GABAergic granule cells (GCs) comprise the largest population of inhibitory neurons (Lepousez et al., 2013; Lledo et al., 2006). A major functional role of GCs is the recurrent and lateral inhibition of output neurons, the mitral and tufted cells (MCs, herein) via dendrodendritic synapses (Isaacson & Strowbridge, 1998; Jahr & Nicoll, 1980). Through these interactions with MCs, GCs are thought to facilitate network oscillations and decorrelation of principal neurons allowing for discrimination of similar odor patterns (Fukunaga et al., 2014; Gschwend et al., 2015; Lepousez & Lledo, 2013; Wanner & Friedrich, 2020). GC activity is regulated by several cell-intrinsic and extrinsinc mechanisms, including excitatory feedback from higher olfactory areas, and several neuromodulatory centers, underscoring their important role in the modulation of MC output and odor processing (Markopolous et al., 2012; Otazu et a., 2015; Boyd et al., 2012; Fletcher & Chen, 2010; Schoppa & Urban, 2003; Schoppa & Westbrook, 1999; Stroh et al., 2012).

Notably, GCs are continuously born throughout life in a process known as adult neurogenesis (Altman, 1962; Altman & Das, 1965). Thus, the dendrodendritic synapses in the OB undergo constant remodeling (Lois & Alvarez-Buylla, 1994; Petreanu & Alvarez-Buylla, 2002). During their integration, adult-born GCs’ (abGCs) dendrites arborize and form functional synaptic contacts with the local circuit and undergo an activity-dependent critical period that affects synaptic connectivity and cell survival (Carleton et al., 2003; Lledo et al., 2006; Petreanu & Alvarez-Buylla, 2002). These changes are accompanied by age-dependent expression of ion channels which ensures their coordinated integration in the OB circuit (Belluzzi et al., 2003; Carleton et al., 2003; Lledo et al., 2006). Although the molecular mechanisms underlying the functional integration of abGCs are not fully understood, this process appears to be highly regulated by both local and afferent synaptic inputs (Lepousez et al., 2015).

A key regulator of both dendrodendritic inhibition (DDI) and the functional integration of abGCs is noradrenaline (NA), released from broadly projecting neurons of the locus coeruleus (Fletcher & Chen, 2010; Jahr & Nicoll, 1982; Moreno et al., 2012; Veyrac et al., 2009). For example, pharmacological blockade of adrenergic receptors in the OB has been shown to impair task-dependent survival of abGCs (Moreno et al., 2012; Veyrac et al., 2009), suggesting that NA affects abGC physiology as early as their critical period of integration. However, the cellular mechanisms underlying the adrenergic regulation of abGC physiology and integration during, and past, the critical period remain poorly understood. NA has been shown to regulate neuronal excitability and network dynamics through a cAMP-dependent hyperpolarization-activated current, I_h_ (Berridge & Waterhouse, 2003; Lüthi A & McCormick D A, 1998; Sara, 2009), through the alteration of intracellular cAMP levels via α_2_- or β-adrenergic receptors (ARs). In previous work, we showed that GCs born just after birth express I_h_, and that this current could influence intrinsic excitability of GCs (Hu et al., 2016). However, the presence of I_h_ in abGCs and its regulation by NA in the OB has not been studied.

Here, using whole-cell patch clamp electrophysiology, we show that several physiological parameters that control neuronal excitability of abGCs show a progressive change during integration, which extends well beyond the critical period. Importantly, I_h_ is present in abGCs and the size of this current increases throughout their maturation. In addition, we show that activation of α_2_-ARs suppresses I_h_ via a cAMP-dependent mechanism. Suppression of I_h_ increases dendritic excitability, thereby enhancing lateral inhibition onto MCs. Furthermore, we show that α_2_-AR activation modulated the I_h_-dependent resonance in GCs. Together these findings suggest that α_2_-AR modulation of I_h_ in abGCs may impart unique features towards fine-tuning interactions of local GABAergic circuits with the OB output neurons in a state-dependent manner.

## METHODS

### Animals

All experiments were conducted following the guidelines of the IACUC of the University of Maryland. Experiments were performed on both adult male and female wild-type (C57/BL6) or Thy1-ChR2 (Jackson Laboratory: stock #007612) mice 2 to 6 months of age, from breeding pairs housed in our animal facility.

### Stereotaxic viral Injections

To label abGCs, mice (P40 – P50) were injected with 200 nL of AAV5-Syn-GFP in the rostral migratory stream (RMS, Addgene), using the following stereotaxic coordinates (in mm): DV −2.85, ML ±0.8, AP +3.3. Anesthetized C57/BL6 mice (2% isoflurane) were head-fixed in a stereotaxic apparatus (Kopf, catalog #940) and a 33-gauge needle (5 uL syringe, Hamilton) was inserted through a 1 mm craniotomy. The speed of virus injection (200 nL/min) was controlled using a syringe pump (Micro4 Microsyringe pump, World Precision Instruments). During the surgery, eyes were lubricated with a petrolatum ophthalmic ointment (Paralube) and body temperature maintained using a heating pad. An intraperitoneal injection of carprofen (5 mg/kg) was used as analgesic.

### Histology and confocal imaging

To characterize the morphological characteristic of abGCs, RMS injected mice were transcardially perfused with cold 4% PFA diluted in 0.1 M PBS, pH 7.4. Brains were then harvested and postfixed overnight at 4°C in the same fixative. Brains were sliced in sections of 100 μm using a vibratome (Leica VT1000). Nuclei were stained with DAPI (Invitrogen) and mounted in a solution of Mowiol-DABCO, prepared as previously described (Villar et al., 2021). Images were acquired using a Leica SP5X confocal microscope (Leica Microsystems) and processed using ImageJ (National Institute of Health).

### Electrophysiology slice preparation

Experiments were performed in OB slices using methods previously described (Hu et al., 2016). Briefly, sagittal or horizontal OB slices were prepared in an oxygenated ice-cold artificial cerebrospinal fluid (ACSF) containing lower Ca^2+^ (0.5 mM) and higher Mg^2+^ (6 mM) compared to normal ACSF. Sections (250 μm) were obtained using a vibratome (Leica VT1000) and then transferred to an incubation chamber containing normal ACSF (see below) and left to recuperate for at least 30 min at 35°C, before the recordings. In all experiments, the extracellular solution is ACSF of the following composition (in mM): 125 NaCl, 26 NaHCO_3_, 1.25 NaH_2_PO_4_, 2.5 KCl, 2 CaCl_2_, 1 MgCl_2_, 1 myo-inositol, 0.4 ascorbic acid, 2 Na-pyruvate, and 15 glucose, continuously oxygenated (95% O_2_-5% CO_2_).

### Recordings and data analysis

Neurons were visualized using an Olympus BX51W1 microscope and recorded using a dual EPC10 amplifier interfaced with the PatchMaster software (HEKA, Harvard Bioscience). Whole-cell recordings were performed at room temperature (∼21°C) or at 32 ± 2°C, using the TC-342B Automatic Temperature Controller (Warner Instruments, Hamden, CT) with a perfusion speed of 2-3 mL/min, using standard patch pipettes (2-8 MΩ resistance). For current-clamp recording, the internal solution had the following composition (in mM): 130 K-gluconate, 4 KCl, 10 HEPES-K, 10 Na phosphocreatine, 2 Na-ATP, 4 Mg-ATP, and 0.3 GTP adjusted to pH 7.3 with KOH (∼290–305 mOsm). For MC voltage-clamp recordings, the K-gluconate was substituted with Cs-gluconate or CsCl. In some of the recordings, the dye Alexa Fluor 594 (20 μM, Invitrogen) was included in the internal recording solution for morphological validation and reconstruction of the recorded neurons.

All electrophysiological recordings were analyzed in MATLAB (Mathworks). Although the whole-cell configuration precludes an accurate measurement of the resting membrane potential (V_m_) due to the cell’s dialysis with the pipette’s internal solution, we estimated V_m_ right after rupturing the seal. Properties of I_h_ were measured as previously described (Hu et al., 2016). Activation kinetics (τ) were determined by fitting a single exponential function to the current trace when stepping from −60 to −130 mV. Voltage dependency was calculated by fitting the tail-current values to a Boltzmann function of the following form: I = I_max_/{1 + exp[(V − V_half_)/k_s_]}, where *I*_*max*_ is the maximal current, *V* is the prepulse potential, *V*_*half*_ is the half-activation potential of the current, and *k*_*s*_ is the slope factor. Statistical significance was determined by Student’s t-test, Kolmogorov– Smirnov test, or one-way ANOVA. Subthreshold resonance was measured using a standard impedance amplitude protocol (ZAP), in which a stimulus of sinusoidal current of constant amplitude and exponentially increasing frequency (0.2 to 20 Hz) is injected into the cell. The ZAP protocol was obtained at −75 mV. The impedance profile was calculated as the magnitude of the ratio of the fast Fourier transforms of the voltage response and current input (Gutfreund et al., 1995; Hutcheon & Yarom, 2000; Vera et al., 2014), using the following equation: Z(f) = |FFT[V(t)]/FFT[I(t)]|. The exponential ZAP protocol we employed works best on cells with lower resonant frequencies (1-4 Hz). The impedance profile was smoothed, and the resonant frequency (f_res_) determined as the frequency at which the maximal impedance value occurred. The strength of resonance (Q factor) was calculated as the ratio between the maximal impedance (|Z(f_res_)|) and the lowest frequency impedance values (|Z(f_low_)|; 0.5 Hz). A neuron was considered resonant if f_res_ > f_low_ and Q > 1.01. Statistical significance was determined by a Student’s t-test.

### Pharmacological agents

Drugs were included in the bathing solution and perfused at a speed of ∼3 mL/min: 4-(N-ethyl-N-phenylamino)-1, 2-dimethyl-6-(methylamino) pyrimidinium chloride (ZD7288), 8-(4-chloro-phenylthio)-2-O-methyladenosine-3,5-cyclic monophosphate (8-CPT-cAMP), N-(2,6-dichlorophenyl)-4,5-dihydro-1H-imidazol-2-amine hydrochloride (clonidine, Clon), 6-cyano-7-nitroquinoxaline-2,3-dione disodium,6-Imino-3-(4-methoxyphenyl)-1(6H)-pyridazinebutanoic acid hydrobromide (gabazine, GBZ), tetrodotoxin (TTX). All drugs were purchased from Tocris (Bio-Techne), except for 8-CPT-cAMP (Abcam).

## RESULTS

### Intrinsic and extrinsic excitability of adult-born granule cells exhibit changes beyond their critical period of integration

To investigate the neurophysiological properties of abGCs during the critical period of their functional integration, we expressed the fluorescent marker GFP in abGCs using viral injections in the RMS of adult mice (see Methods). We recorded abGCs at two time points: 2- and 6-weeks post-injection (2 wk and 6 wk old, respectively). Previous studies have shown that during this time window abGCs make functional connections with the pre-existing circuit; at 2 wk, GCs are undergoing a critical period of synaptic remodeling, whereas cells at 6 weeks are considered to be fully integrated (Lepousez et al., 2013; Lledo et al., 2006; Petreanu & Alvarez-Buylla, 2002). Two weeks post-injection, abGCs exhibit mature morphological characteristics, including a primary apical dendrite, along with distal branches and prominent spines, with 53% of abGCs corresponding to class V, 37% to class IV, and 10% to class III, as previously described (class V, 41/78; class IV, 29/78; class III, 8/78; n = 3 mice; **Figure 1A**) (Carleton et al., 2003; Petreanu & Alvarez-Buylla, 2002). Interestingly, despite the degree of maturation exhibited by abGCs at 2 wk they continued to exhibit significant changes in excitability as they futher matured. Specifically, the resting membrane potential (RMP) and input resistance were significantly different between these two age groups. At 2 wk, abGCs had a more depolarized RMP than at 6 wk (–67.9 ± 1.2 mV, 2 wk, n = 16 cells; –73.1 ± 1.1 mV, 6 wk; n = 18 cells; p = 0.003; **Figure 1A**) and had a higher input resistance (1.64 ± 0.24 GΩ vs. 0.56 ± 0.06 GΩ; p = 0.0001). In contrast, we did not find differences in cell capacitance (35.7 ± 3.7 pF vs. 36.2 ± 2.8 pF; p = 0.91). However, even though abGCs at 6 wk exhibited a hyperpolarized RMP and lower input resistance they had a higher maximum frequency of firing in response to current stimuli (2 wk = 17.8 ± 2.3 Hz; 6 wk = 32.3 ± 2.8 Hz; p = 0.0005). Accordingly, a linear fitting to the input-output function revealed a shallower slope for older abGCs (2 wk = 0.73 ± 0.09 Hz/pA; 6 wk = 0.29 ± 0.03 Hz/pA; p = 0.00003; **Figure 1C**), suggesting a change in gain function as abGCs mature. In addition to changes in excitability due to intrinsic membrane properties, there were marked changes in the spontaneous excitatory postsynaptic currents (sEPSCs) as the abGC matured (**Supplementary Figure 1**). The frequency of sEPSCs increased by two-fold with the maturation of the abGCs (2 wk = 2.8 ± 0.7 Hz; 6 wk = 6.8 ± 1.5 Hz; p = 0.03; **Supplementary Figure 1 A, B**), together with an increase in the average amplitude of the sEPSCs (2 wk = 10.1 ± 0.5 pA, 6 wk = 13.9 ± 1.3 pA; p = 0.02). Lastly, both the rise and decay times of the EPSCs decreased with cell-age (rise time, 2 wk = 1.7 ± 0.1 ms, 6 wk = 1.1 ± 0.08 ms; p = 0.0001; decay, 2 wk = 6.8 ± 0.3 ms, 6 wk = 5.1 ± 0.4 ms; p = 0.003). Together, these results indicate that as abGCs functionally integrate, between 2 and 6 wk of cell age, the apparent decrease in excitability is offset by a large increase in the number of excitatory inputs, which exhibit distinct kinetic properties.

**Figure 1.**
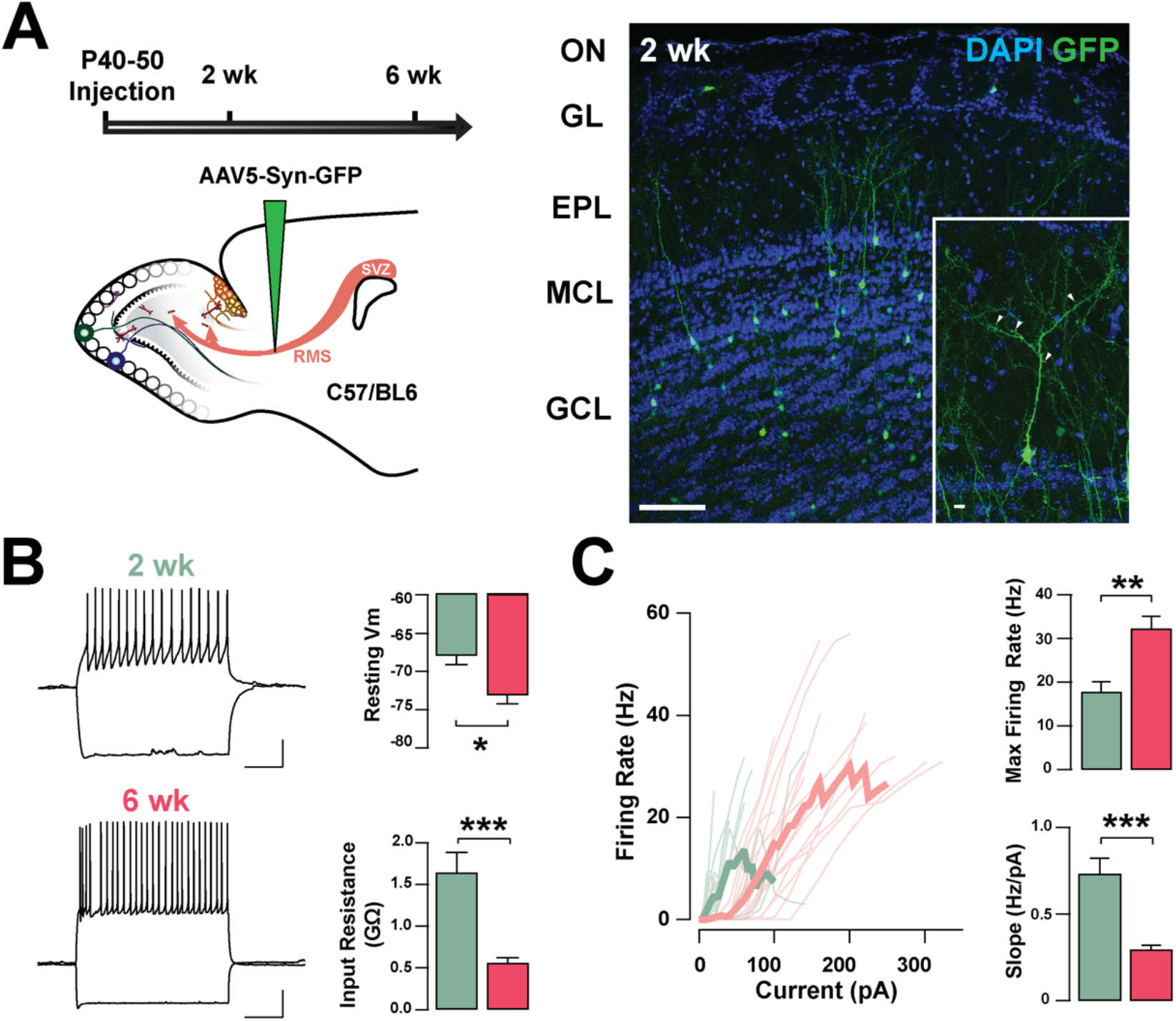
Intrinsic physiological properties of adult-born GCs **(A)** Left: diagram of strategy used for labeling abGCs. Adult-born neurons were labeled with GFP at 40-50 days postnatal (p40-50) using virus-mediated transduction. We examined GC properties at 2- and 6-weeks (wk) post-injection. Right: Confocal images of GFP-labeled adult-born neurons (green) in the MOB at 2 wk post-injection. The inset shows a mature GC with a primary apical dendritic branch and abundant dendritic spines (arrow heads). OB cell layers, indicated on the left, were determined using DAPI (blue). Calibration: 100 μm and 10 μm (inset). **(B)** Left: traces from representative cells at 2 wk (green) and 6 wk (pink) post-injection showing responses to square current steps (–120 pA to +260 pA). The resting membrane potential (Vm) was –63 mV and –78 mV, respectively. The voltage traces show sustained action potentials in response to positive current injections and a small sag with negative current injections. The calibration is 20 mV and 500 ms. Right: bar graphs summarizing changes in GC excitability at two stages of cell maturation (2 wk, n = 16 cells; 6 wk, n = 18 cells). From 2 to 6 wk post-injection, abGCs show a decrease in resting Vm (p = 0.003), and a decrease in the input resistance (p = 0.0001). **(C)** Left: input-output curves for firing rates elicited by a range of 2 s square current pulses. Right: Cells at 6 wk post-labeling exhibited a higher maximal firing rate (p = 0.0005), and lesser slope in their input-output function (p = 0.00003).

### Adult-born granule cells exhibit I_h_

Previously, we described the presence of I_h_ in GCs labeled born just after birth (P1-P6), and found that this current increased in amplitude during postnatal maturation (Hu et al., 2016); therefore, we examined the presence of I_h_ in abGCs. As shown in Figure 2A, stepping to negative potentials (–60 to –130 mV), revealed a characteristic inward current at both 2 and 6 wk; however, at –130 mV the I_h_ was ∼40% greater at 6 wk (2 wk = –72.9 ± 9.6 pA, n = 17 cells; 6 wk = –101.4 ± 9.4, n = 18 cells; p = 0.03). Because the subunit composition of hyperpolarization-activated cyclic nucleotide-gated (HCN) channels determines the kinetic properties and voltage dependency of the currents (Shah, 2014; Zagotta et al., 2003), we examined the activation kinetics and the voltage-dependency of I_h_ at both ages. The rate of activation and the voltage-dependency were not different at these two time points (τ; 2 wk = 57.3 ± 11.1 ms, 6 wk = 53.7 ± 9.6 ms, p = 0.8; V_half_: 2 wk = –98.8 ± 1.2 mV, 6 wk = –100.2 ± 2.2 mV; p = 0.6; **Figure 2B**). These results indicate that abGCs, like postnatal born GCs (Hu et al., 2016), exhibit an age-dependent increase in I_h_, with no change in the kinetic properties.

**Figure 2.**
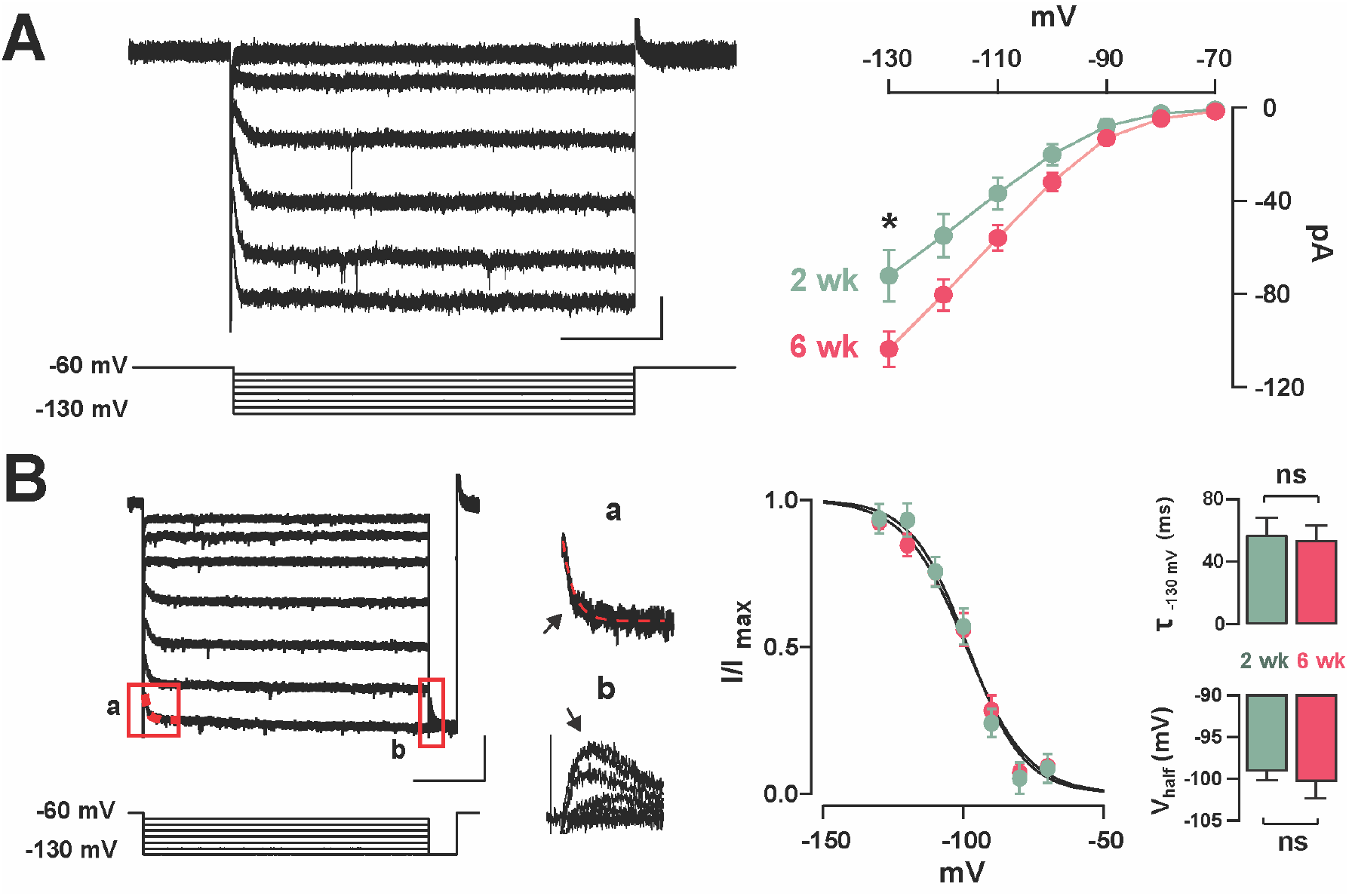
Functional characteristics of I_h_ in adult-born GCs **(A)** Left: hyperpolarizing voltage steps from −60 to −130 mV (10 mV increments) elicited a hyperpolarization-activated inward current (I_h_) in a GC 2 wk post-labeling. The calibration is 20 pA and 500 ms. Right: The current–voltage (I–V) relationship for I_h_ in adult born GCs at 2 and 6 wk post-injection shows an increase in I_h_ amplitude with cell age (2 wk, n = 17 cells; 6 wk, n = 18 cells; current at –130 mV, p = 0.03). **(B)** Left: hyperpolarizing steps from −60 to−130 mV with a test pulse to −130 mV were used to elicit a tail current. The calibration is 50 pA and 500 ms. Expanded currents traces show examples for a single exponential fit for estimating activation kinetics (a; τ = 43 ms, red dotted line), and for the tail current amplitude to determine voltage dependency (b). Right: Boltzmann fits for I_h_ in GCs at 2 and 6 wk post-labeling show no difference in the activation curve for I_h_. Summary bar graphs showing the similar average for the activation kinetic (upper right, τ_130mV_, p = 0.8) and half activation potential (bottom right, Vhalf, p = 0.6) for the two ages, post-labeling.

### Adrenergic modulation of I_h_ in adult-born GCs

Previous studies have shown that NA can regulate I_h_ through the activation of α_2_-AR, which through a G_i_-coupled pathway, reduces cAMP and leads to a reduction in I_h_ (Barth et al., 2008; Umemura et al., 1986). To examine the possibility that NA acts directly in abGCs to regulate I_h_, as opposed to a disynaptic effect, we conducted experiments in the presence of blockers of synaptic transmission and voltage-dependent sodium channels (10 μM CNQX, 50 μM APV, 10 μM GBZ, 0.5 μM TTX). Under these conditions, NA reduced I_h_ by approximately 20% the normalized current (at –130 mV), in both 2 and 6 wk old abGCs (2 wk = 0.77 ± 0.06, n = 5 cells, p = 0.07; 6 wk = 0.82 ± 0.03; n = 4 cells; p = 0.02; **Figure 3A**). Importantly, application of the selective α_2_-AR agonist, clonidine (Clon, 10 μM), similarly reduced I_h_ in abGCs, at both 2 and 6 wk (2 wk = 0.79 ± 0.06, n = 6 cells, p = 0.03; 6 wk = 0.77 ± 0.04; n = 7 cells; p = 0.001; **Figure 2A**). To examine the possibility that the reduction in I_h_ by α_2_-AR activation is due to changes in intracellular cAMP, we conducted additional pharmacological experiments in 6 wk abGCs. Application of the membrane-permeable cAMP analogue, 8-CPT-cAMP (8-CPT, 100 μM), increased I_h_ by 15% (normalized current, 1.15 ± 0.03; n = 5 cells; p = 0.01; **Figure 3A**), while pre-application of 8-CPT occluded the Clon-mediated suppression of I_h_ (normalized current, 0.93 ± 0.09; n = 5 cells; p = 0.45). In agreement with a regulation of the HCN by cAMP, Clon shifted the voltage-current relationship of I_h_ to more negative potentials (V_half_: Control = –98 ± 3.4 mV; Clon = –105.5 ± 3.4 mV, p = 0.04; **Figure 3B**), while the addition of 8-CPT shifted the V_half_ to more positive potentials (–94.4 ± 3.0 ms, p = 0.02). In contrast, the rate of activation of I_h_ at –130 mV did not change by α_2_-AR activation or by the addition of 8-CPT (Control, CTL = 53.7 ± 9.5 ms; Clon = 47.7 ± 12.5 ms, p = 0.16; 8-CPT = 49.7 ± 14.2 ms; p = 0.54; **Figure 3B**). However, pre-application of the cAMP analog occluded the change in voltage dependency elicited by Clon (8CPT + Clon = –99.1 ± 4.2 mV, p = 0.58, not shown). Lastly, in agreement with the decrease in I_h_ elicited by α_2_-AR activation, Clon also produced an apparent increase in input resistance at both time points, albeit the effect was only significant at 6 wk (fold-increase: 2 wk = 1.29 ± 0.1, p = 0.08; 6 wk = 1.37 ± 0.08, p = 0.01; not shown). Together, these results indicate that I_h_ is present as early as the critical period during the functional integration of abGCs. Importantly, NA acting through α_2_-ARs can modulate I_h_ in abGCs indicating another cellular mechanism by which NA can regulate the intrinsic excitabity of GCs in the OB (Smith et al., 2009; Zimnik et al., 2013).

**Figure 3.**
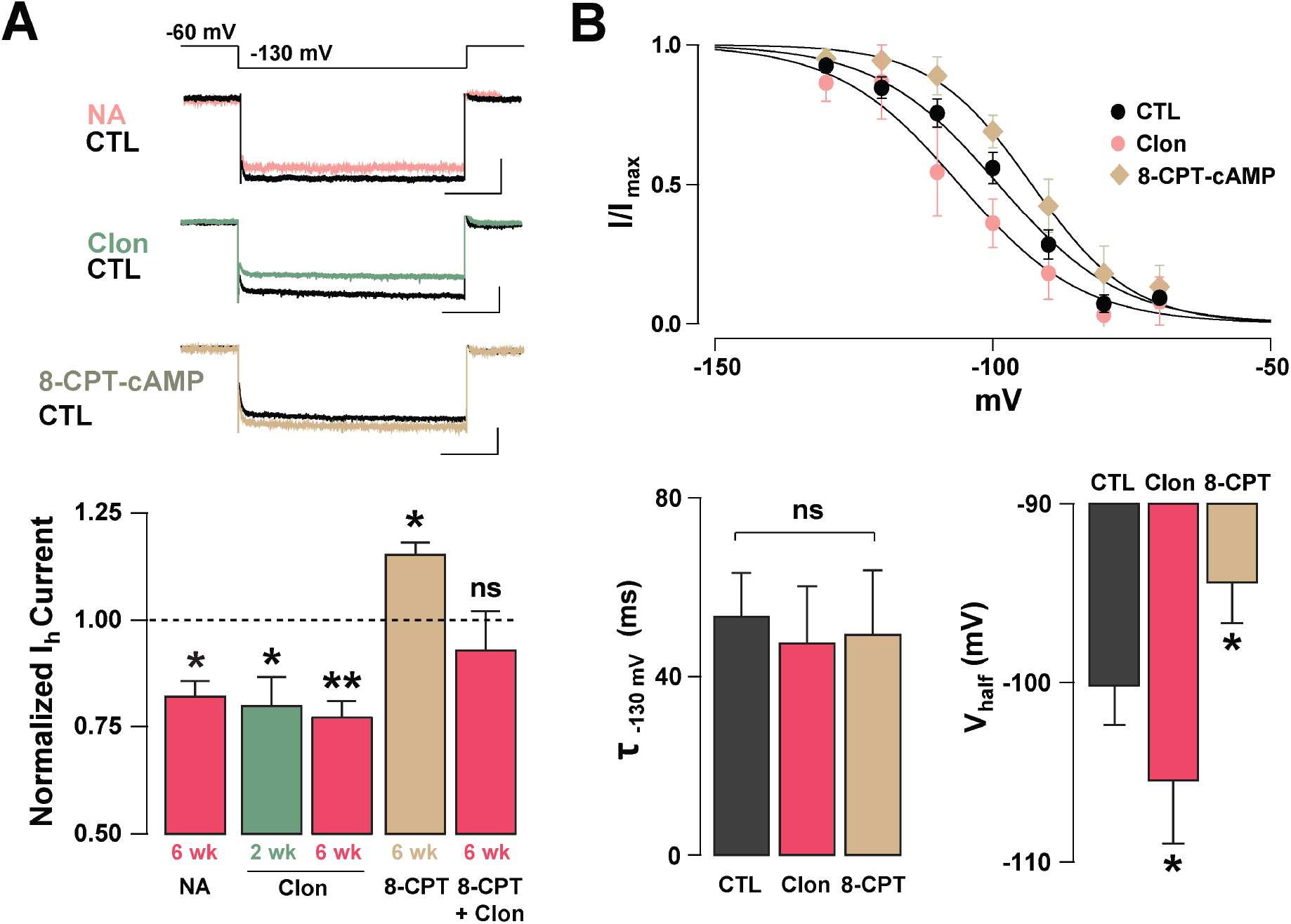
NA reduces I_h_ in adult-born GCs via activation of α2-ARs **(A)** Top: example traces of I_h_ elicited with a hyperpolarizing step from −60 to −130 mV, control (CTL) and in the presence of drugs. NA reduced the I_h_ in a GC 6 wk post-labeling (CTL, black; NA 10 μM, pink). A similar reduction is elicited by the selective α_2_-AR agonist, clonidine (Clon, 10 μM, green). In contrast, application of a membrane-permeable cAMP analog, 8-CPT-cAMP (100 μM, tan) produced an increase in I_h_. The calibration is 50 pA and 500 ms. Bottom: Summary bar graphs for the effect of the pharmacological treatments on the normalized I_h_ amplitude at −130 mV, with respect to control. NA and Clon reduced the I_h_ amplitude in GCs (NA, 6 wk, n = 4 cells, p = 0.02; Clon, 2 wk, n = 6 cells, p = 0.03; Clon 6 wk, n = 7 cells, p = 0.001), while I_h_ amplitude is increased by the addition of 8-CPT-cAMP (n = 5 cells, p = 0.01). In the presence of 8-CPT-cAMP plus Clon, the I_h_ amplitude was not significantly different from CTL (n = 5 cells, p = 0.45). **(B)** Top: boltzmann fits for the voltage-dependency of activation of I_h_ in GCs 6 wk post-injection showing that activation of α_2_-ARs by Clon and application of 8-CPT-cAMP shifts the voltage-dependency of I_h_ in opposite directions. Bottom: summary bar graphs showing that application of Clon or 8-CPT-cAMP does not alter the activation kinetics of I_h_ (Clon, p = 0.16; 8CPT, p = 0.54), but that α2-AR activation shifts the half-activation potential (V_half_) to more negative potentials (p = 0.04), whereas 8-CPT-cAMP shifts the V_half_ to more positive potentials (p = 0.02).

### I_h_-mediated resonance in GCs is modulated by α_2_-AR activation

We have previously shown that subthreshold resonance in the theta (θ) frequency range in postnatally born GCs relies on I_h_ activation (Hu et al., 2016). Thus, we hypothesized that modulation of I_h_ by α_2_-AR activation could regulate subthreshold resonance in mature GCs. A sinusoidal current stimulus of increasing frequency and constant amplitude injected into GCs (ZAP; see Methods) revealed resonance in a subset of GCs in adult mice (**Figure 4A**). At –75 mV, the resonance strength (Q-factor) was 1.18 ± 0.06 and the resonant frequency (F_res_) was 1.98 ± 0.28 Hz, consistent with data previously taken at similar membrane potentials in postnatal GCs (Hu et al., 2016). Clon produced a noticeable decrease in both the Q-factor and the F_res_ (1.05 ± 0.02; p<0.02 and 1.00 ± 0.23 Hz; n = 6 cells; p < 0.0002; **Figure 4B**). Furthermore, Clon increased the input resistance of GCs, in a manner consistent with a reduction in I_h_ (control, 1.15 ± 0.16 GΩ vs. Clon, 2.31 ± 0.46 GΩ; n = 6 cells; p < 0.02, not shown). Additionally, 8-CPT-cAMP occluded the effect of Clon on both F_res_ and Q-factor (f_res_, 8-CPT-cAMP, 1.10 ± 0.24 Hz vs. 8-CPT-cAMP + Clon, 1.01 ± 0.32 Hz, p=0.82; Q-factor, 8-CPT-cAMP, 1.07 ± 0.03 vs. 8-CPT-cAMP + Clon, 1.03 ± 0.02, p=0.2; n=3 cells; not shown). These results indicate that activation of α_2_-ARs by NA modulates the resonant properties of GC in a manner consistent with a reduction of intracellular cAMP.

**Figure 4.**
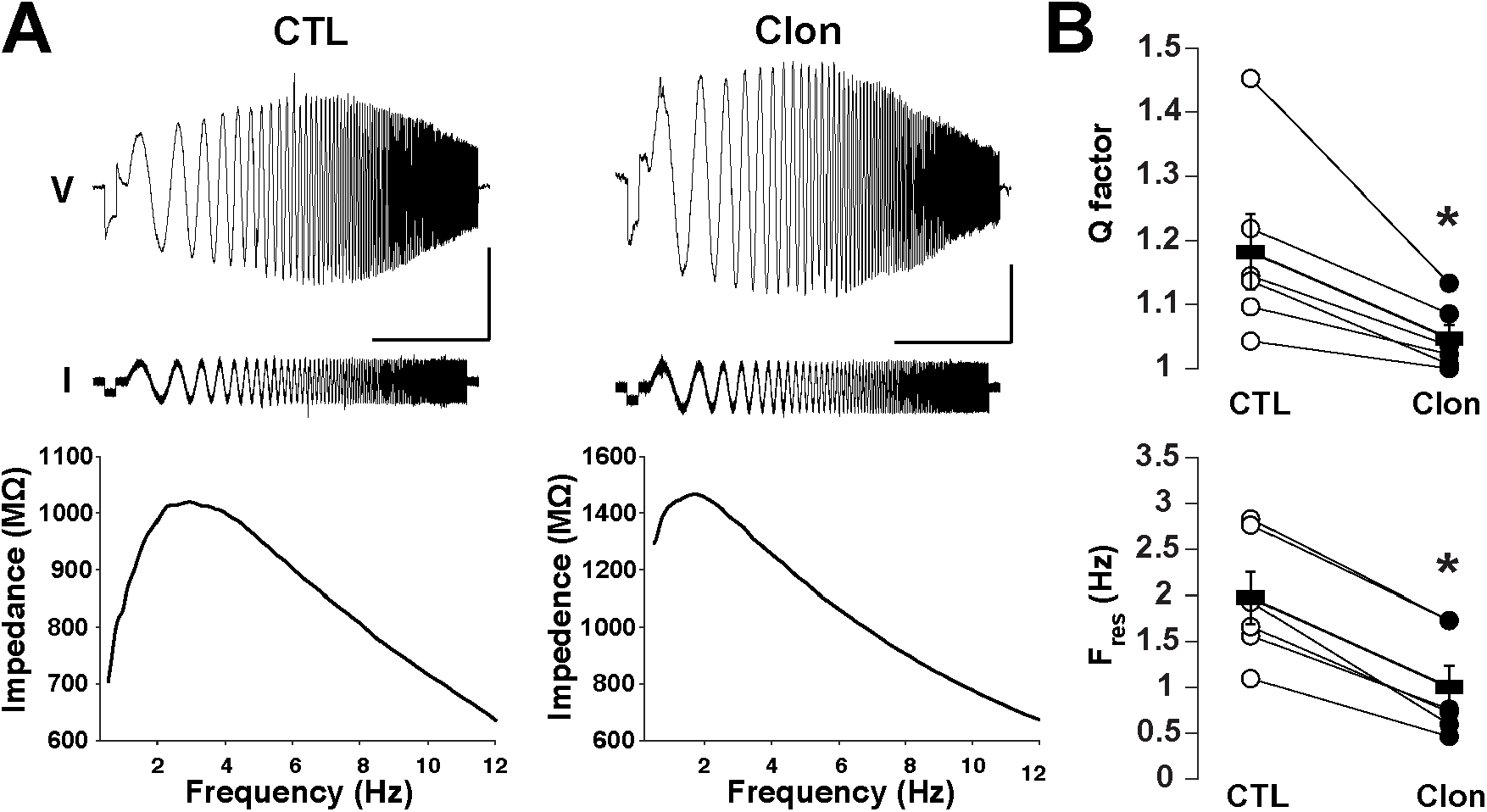
Activation of α_2_-ARs decreases the resonance of GCs **(A)** Top: Voltage responses (upper traces) to a ZAP current stimulus (lower traces) in a GC before (left) and after the application of Clon (10 μM, right). The calibration is 5 mV and 10 s; the ZAP stimulus amplitude is 5 pA (peak-to-peak). Bottom: impedance profiles for the trials shown on the figure above. In the control condition, the resonant strength (Q factor) is 1.45 and the peak frequency 2.83 Hz. In the presence of Clon (right), the resonant strength is 1.13 and the peak frequency is 1.73 Hz (right). The Vm was maintained at –75 mV for both trials. **(B)** Summary plots for the effect of Clon. Activation of α_2_-ARs by the agonist significantly reduced both Q factor and resonant frequency (F_res_) in GCs.

### α_2_-AR activation modulates dendritic excitability in abGCs and reciprocal dendrodendritic inhibition

We hypothesize that I_h_ is a target for the regulation of dendritic excitability in GCs and therefore the modulation of DDI at GC-MC synapses. To this extent, we recorded GABAergic currents from MCs in symmetric chloride conditions and elicited DDI by briefly (50 ms) depolarizing MCs (–70 to 0 mV). To isolate the dendritic output from GCs, we performed these experiments in the presence of TTX (0.5 μM) and in 0 Mg^2+^. Under these conditions, GABA release from GC dendrites is solely driven by the activation of both AMPA and NMDA receptors upon release of glutamate from MCs. A depolarizing step (50 ms) elicited a large inward current, with a slow decay (369 ± 43 ms, n = 26 cells), similar to the time course of DDI previously reported by us and others (Isaacson & Strowbridge, 1998; Schoppa et al., 1998; Villar et al., 2021). This current was completely blocked by the GABA_A_ receptor antagonist, gabazine (GBZ, 10 μM; n = 4 cells; **Figure 5A**). Quantification of the transferred charge indicated that Clon produced a significant increase in DDI (CTL, –110.7 ± 17.5 pC, Clon, –157.6 ± 25.6 pC; n = 10 cells; p = 0.01; **Figure 5B**), with a slight increase, although not significant, in the rate of decay of the DDI current (CTL, 315 ± 57 ms, Clon, 479 ± 53; p = 0.08; not shown). Importantly, the selective I_h_ blocker ZD7288 (ZD, 10 μM) also increased DDI onto MCs (CTL, –83.1 ± 15.8 pC, ZD, –189.2 ± 37.7 pC, n = 16 cells, p = 0.003, **Figure 5B**). Furthermore, pre-application of ZD occluded the increase in DDI by α_2_-AR activation (ZD, –115.3 ± 22.6 pC, ZD + Clon, –138.4 ± 32.4 pC; n = 7 cells; p = 0.14; **Figure 5B**), suggesting that the α_2_-mediated increase in DDI occurs via a suppression of I_h_. Surprisingly, however, when we used a physiological concentration of external Mg^2+^ (1 mM), α_2_-AR activation resulted in a decrease in DDI (CTL, –83.4 ± 9.4 pC, Clon, –60.2 ± 9.4, n = 15 cells, p = 0.006, **Figure 5C**); similarly, application of the I_h_ blocker also resulted in a small reduction in DDI (CTL, –66.4 ± 8.2 pC, ZD, –51.3 ± 5.0 pC; n = 13 cells; p = 0.03, **Figure 5C**). We reason that it is possible that under physiological blockade of NMDA receptors by Mg^2+^, a suppression of I_h_ by α_2_-AR activation could hyperpolarize the dendrites of GCs, as we previously showed (Hu et al., 2016), thus reducing the release of GABA from GCs.

**Figure 5.**
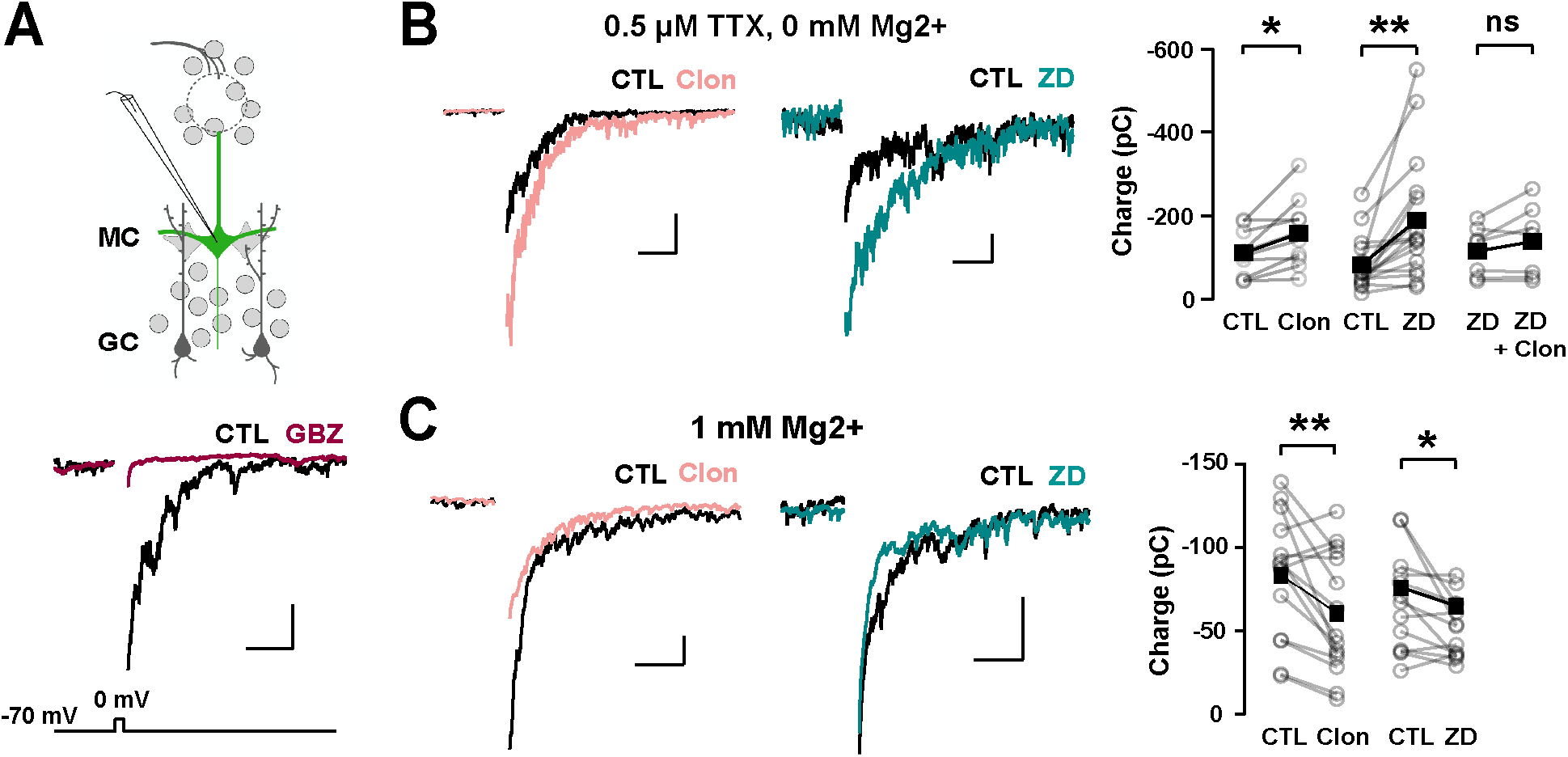
α_2_-AR modulation of I_h_ on recurrent inhibition and GC dendritic excitability **(A)** Top: recording scheme for eliciting reciprocal dendritic inhibition. We recorded recurrent inhibition elicited by a depolarizing voltage step in MCs in slices from the Thy1-ChR2 mouse. Sample traces show dendrodendritic inhibition, in which MCs were depolarized from –70 to 0 mV to recruit recurrent inhibition. Bottom: the current is abolished by the GABA_A_ antagonist gabazine (GBZ, 10 μM). **(B)** Representative averaged traces of recurrent inhibiton obtained in TTX (0.5 μM) and Mg^2+^-free ACSF, and summary plots. Activation of α_2_-ARs by Clon (10 μM; pink, n = 10 cells, p = 0.01) and blockade of I_h_ with ZD7288 (ZD,10 μM; teal, n = 16 cells, p = 0.003) increased the recruited inhibition. Clon failed to increase the recruited inhibition with the pre-application of ZD (n = 7 cells, p = 0.14). The calibration is 100 pA and 500 ms. **(C)** In physiological ACSF (1 mM Mg^2+^ and no TTX), α_2_-AR activation by Clon and I_h_ blockade by ZD failed to increase recurrent inhibition in MCs. Instead, a small reduction is observed in the presence of ZD (Clon, n = 15, p = 0.006; ZD, n = 13 cells, p = 0.03). The calibration is 50 pA, 200 ms.

### α2-AR suppression of I_h_ enhances lateral inhibition in the MOB

In addition to directly modulating recurrent inhibition at a given GC-MC synaptic pair, changes in GC dendritic excitability by I_h_ could alter lateral inhibition between MCs. To test this possibility, we elicited lateral inhibition in MCs using the Thy1-ChR2 mouse, in which the light-gated cation channel channelrhodopsin (ChR2) is expressed in MCs under the *Thy1* promoter (Arenkiel et al., 2007). In cell-attached configuration, we adjusted the light intensity so that a 2 ms light pulse reliably produced a single action potential in the MCs, with a short latency (under 5 ms, **Figure 6A**). We then recorded in voltage-clamp using a Cs-gluconate based internal solution to isolate disynaptic lateral inhibition in the recorded MC from light-induced feedforward excitation. At –70 mV, light activation elicited a fast inward current (onset 6.3 ± 0.2 ms, not shown), while at the reversal potential for excitatory currents (∼ +10 mV under the recording conditions), light stimulation produced a large outward current (249 ± 28 pA, n = 41 cells), that occurred with a longer delay (9.5 ± 0.2 ms, **Figure 6A**). In physiological Mg^2+^, the outward current had a faster decay (90.8 ± 12.9 ms) and was completely abolished by GBZ (10 μM; n = 3 cells; **Figure 6A**) consistent with the activation of dendrodendritic synapses from neighboring MCs. We note that at depolarized potentials (+10 mV), voltage-gated sodium and calcium channels are likely inactivated and therefore the outward current elicited by light stimulation is unlikely to result from recurrent inhibition from the recorded MCs. Therefore, the light-evoked disynaptic inhibition is likely to contain a predominant component from lateral inhibition.

**Figure 6.**
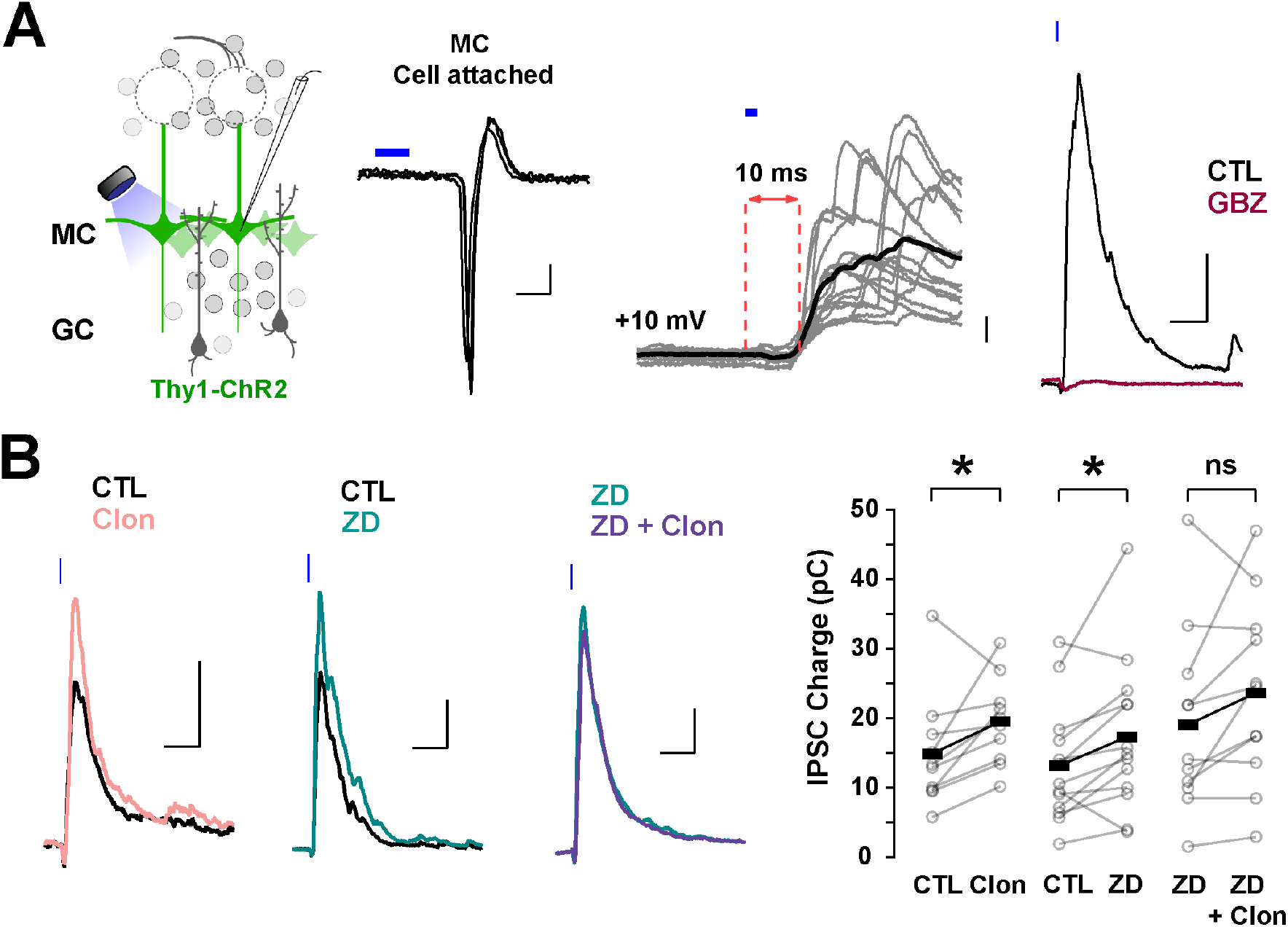
Suppression of I_h_ by α2-AR increases lateral inhibition onto MCs **(A)** Left: diagram of the strategy used to elicit lateral inhibition. We recorded from and optogenetically-activated MCs expressing the ChR2 protein (Thy1-ChR2 mouse). Right: example of a cell-attached recording showing that activation of ChR2 (2 ms) elicits direct, reliable action potentials in the recorded MC. In voltage-clamp recordings at +10 mV, ChR2 stimulation evokes a disynaptic inhibitory current (10 ms delay) which is abolished by GBZ (10 μM, brown). The calibration is: cell-attached, 20 pA, 2 ms; whole cell, 50 pA, 50 ms. **(B)** Representative averaged traces (left) and summary plots (right) showing that α2-AR activation by Clon and blockade of I h increased the light-evoked inhibitory currents (Clon, pink, n = 10, p = 0.04; ZD, teal, n = 13 cells, p = 0.02). In the presence of ZD, Clon failed to increase the light-evoked inhibitory currents (purple; n = 11 cells, p = 0.09).

Using this protocol to elicit lateral inhibition, we found that Clon produced an increase in the peak outward current (CTL, 212.1 ± 57.2 pA, Clon, 244.2 ± 61.5 pA; n = 10 cells; p = 0.002; **Figure 6B**), and the charge (CTL, 14.9 ± 2.6 pC, Clon,19.5 ± 1.9 pC, p= 0.04), without affecting the rate of decay of the outward current (CTL, 141± 47 ms, vs Clon, 99 ± 24 ms, p = 0.44). Similarly, blockade of I_h_ with ZD increased both the peak current and the transferred charge (CTL, 245.2 ± 39.7 pA, ZD, 304.6 ± 53.3 pA; p = 0.002; CTL, 13.2 ± 2.3 pC, ZD, 17.3 ± 3.1 pC; p = 0.02; n = 13 cells; **Figure 6B**). As with the recurrent inhibition in 0 mM external Mg^2+^, pre-application of ZD occluded the increase in lateral inhibition elicited by α_2_-AR activation (ZD, 313.7 ± 64.7 pA, ZD + Clo, 297.7 ± 64.4 pA; p = 0.30; ZD, 19.2 ± 4.0 pC, ZD + Clon, 23.7 ± 4.0 pC, p = 0.09; n = 11 cells; **Figure 6B**). These results indicate that suppression of I_h_ by α_2_-AR activation can increase the amplitude of lateral inhibition, similar to the observed increase in GC dendritic output in the experiments described above in TTX and Mg^2+^-free (**Figure 5B**).

The increase in lateral inhibition in MCs by α_2_-AR activation likely results from an increase in inhibition from GC to MC rather than an increase in excitatory transmission MC-GC, as Clon did not affect the evoked EPSCs on GCs, recorded at –70 mV (peak current, CTL, –33.1 ± 7.7 pA, Clon, –25.7 ± 6.1 pA; p = 0.39; charge, CTL, –1.12 ± 0.20 pC, Clon, –0.91 ± 0.28 pC; p = 0.39; n = 5 cells; **Supplementary Figure 2**). A similar result was obtained with ZD (peak current, CTL, –50.6 ± 14.6 pA, ZD, –56.9 ± 20.2 pA, p = 0.59; charge, CTL, –1.69 ± 0.62 pC, ZD, –1.76 ± 0.62 pC; p =0.73; n = 5 cells). In summary, these results indicate that modulation of I_h_ by α_2_-AR modulation results from an increase in transmission at GC-MC synapses.

## DISCUSSION

The control of dendritic excitability in GCs is crucial for regulating dendrodendritic interactions with MCs, with both reciprocal and lateral inhibition affecting the spatiotemporal decorrelation of MC spiking activity in the OB (Gschwend et al., 2015; Wanner & Friedrich, 2020; Wiechert et al., 2010). These forms of inhibition are regulated by state-dependent noradrenergic modulation from the locus coeruleus (LC); however, beyond the description of changes in excitability in some cell types of the OB, the mechanisms of this regulation are poorly understood (Jahr & Nicoll, 1982; Nai et al., 2009; Shipley et al., 1985; Trombley & Shepherd, 1992; Zimnik et al., 2013). Furthermore, inhibitory circuits within the OB undergo an exceptional remodeling throughout life due to the neurogenesis of GCs through a highly regulated process (Gheusi & Lledo, 2014; Lepousez et al., 2013; Lledo & Saghatelyan, 2005), including regulation by NA (Fletcher & Chen, 2010; Jahr & Nicoll, 1982; Moreno et al., 2012; Veyrac et al., 2009). Here, we provide evidence that NA, acting on α_2_-ARs, can regulate the excitability of abGCs by reducing I_h_ via a cAMP-dependent mechanism. This suppression of I_h_ increases dendritic excitability, thereby enhancing lateral inhibition onto MCs and modulates the resonance of GCs. Together, these findings provide a novel mechanism by which adrenergic modulation of intrinsic properties of abGCs can regulate olfactory function in a state-dependent manner.

While prior studies have examined the progression of intrinsic and synaptic properties of abGCs in the OB as they matured, their primary focus was on earlier developmental time points leading up to 2 to 3 weeks of cell age (Belluzzi et al., 2003; Carleton et al., 2003). Here, we show that abGCs undergo further electrophysiological changes past this critical period with changes in basic electrophysiological parameters that result in a general decrease in intrinsic excitability between 3 to 6 weeks of age. These changes can provide a mechanism for the observed changes in odor tuning and response profiles in abGCs as a function of maturation (Magavi et al., 2005; Wallace et al., 2017). In addtion, the increase in synaptic activity we observed, as abGCs matured beyond 2 wk, aligns with previous studies showing an increase in the number of MC-GC synapses during development and changes in the relative expression of glutamatergic receptors such AMPA and NMDA receptors as GC mature (Carleton et al., 2003; Dietz et al., 2011; Hinds & Hinds, 1976). Interestingly, the changes in intrinsic excitability, in particular the expression of I_h_, mirrored those of postnatally born neurons previously described (Hu et al., 2016), suggesting that in terms of I_h_ expression, adult neurogenesis recapitulates developmental or postnatal neurogenesis.

The voltage dependency and activation kinetics of HCN channels are modulated by intracellular levels of cAMP, implicating I_h_ in the regulation of state-dependent network activity by neuromodulators that regulate cAMP concentrations such as NA (He et al., 2014; Lüthi A & McCormick D A, 1998; McCormick et al., 1991; Surges et al., 2006). Our studies indicate the presence of α_2_-ARs in abGCs and that their activation produced modulation of I_h_, as early as 2 weeks post-labeling in the RMS. Furthermore, abGCs during their functional stages of integration exhibit an α_2_-AR-mediated reduction of I_h_, by shifting the voltage dependency to lower membrane potentials. This agrees with our results showing that bath application of 8-CPT occluded the α_2_-mediated suppression of I_h_ relative to control conditions, as well as shifting its voltage-dependency in abGCs.

α_2_-mediated changes in I_h_ also reduced resonance in GCs. While it is unclear the effect that this decrease in resonance will have on odor processing in the OB, inputs into the MOB are driven by respiration, which occurs in the θ frequency domain, potentially allowing increased excitation of GCs at θ frequencies through this bandpass mechanism, thus altering the timing of GC to MC inhibition. Furthermore, the reduction in the resonance of GCs by α_2_-AR activation may affect the temporal integration of excitatory inputs into the dendrites of GC. Studies in the hippocampus have shown that I_h_ can act as a bandpass filter, allowing synaptic inputs at θ frequencies to be preferentially transferred to the soma (Narayanan & Johnston, 2008).

Our experimental paradigm allowed us to examine the neuromodulation by NA of reciprocal and lateral inhibition in isolation. Previous studies of dendrodendritic inhibition have often used MC depolarization to elicit recurrent inhibition (Isaacson, 2001; Isaacson & Strowbridge, 1998), or glomerular stimulation (Pressler et al., 2007; Schoppa, 2006; Schoppa et al., 1998), which fails to distinguish the contribution of reciprocal and lateral inhibition. Surprisingly, the effect of NA in reciprocal inhibition was Mg^2+^-sensitive. In the absence of Mg^2+^, α_2_-AR activation and blockade of I_h_ increased the GABAergic output from GCs. However, in the presence of physiological Mg^2+^, we observed the opposite effect. We hypothesize that the α_2_-AR suppression of I_h_ would not only increase spine or dendrite resistance but could also hyperpolarize these compartments (Hu et al., 2016). In agreement, a similar effect has been observed in pre-synaptic compartments such as the calyx of Held (Huang & Trussell, 2014). At the level of the GC dendritic arbors or a dendritic spine, the increased resistance from the suppression of I_h_ allows for larger changes in voltage with glutamate receptor activation. This agrees with other studies showing an increase in dendritic excitability with α_2_-AR suppression of I_h_ in pyramidal neurons (Barth et al., 2007, 2008; Labarrera et al., 2018; Wang et al., 2007). However, in GCs, the concurrent hyperpolarization induced by suppression of I_h_ would also shift the membrane potential farther from the voltage threshold for the release of the Mg^2+^ block in the NMDA receptors, as the α_2_-mediated increase in reciprocal inhibition was observed only in the absence of Mg^2+^.

Taken together, our results show that NA can modulate inhibitory functions of GCs via the regulation of I_h_. At a circuit level, the enhancement of lateral, but not reciprocal inhibition, by α_2_-ARs may coordinate MC output to enhance separation of MC activity to allow for the narrowing of odor tuning curves in MCs (Yokoi et al., 1995). Due to the observed physiological differences between young and old abGCs, future studies will address the neuromodulation of dendritic computation in an age-dependent manner.

## ACKNOWLEDGMENTS

This research was supported by National Institutes of Health (NIH)-National Institute on Aging Grant AG-049937A (R.C.A) and National Science Foundation-Graduate Research Fellowships Program/Division of Graduate Education Grant 1322106 (R.H.).

## AUTHOR CONTRIBUTIONS

R.H and R.C.A. designed research; R.H., J.S, G.Z.D. and P.S.V. performed research; R.H., J.S, G.Z.D. and P.S.V. analyzed data; R.H., J.S., P.S.V. and R.C.A. wrote the paper.

## FIGURES AND LEGENDS

**Supplementary Figure 1.**
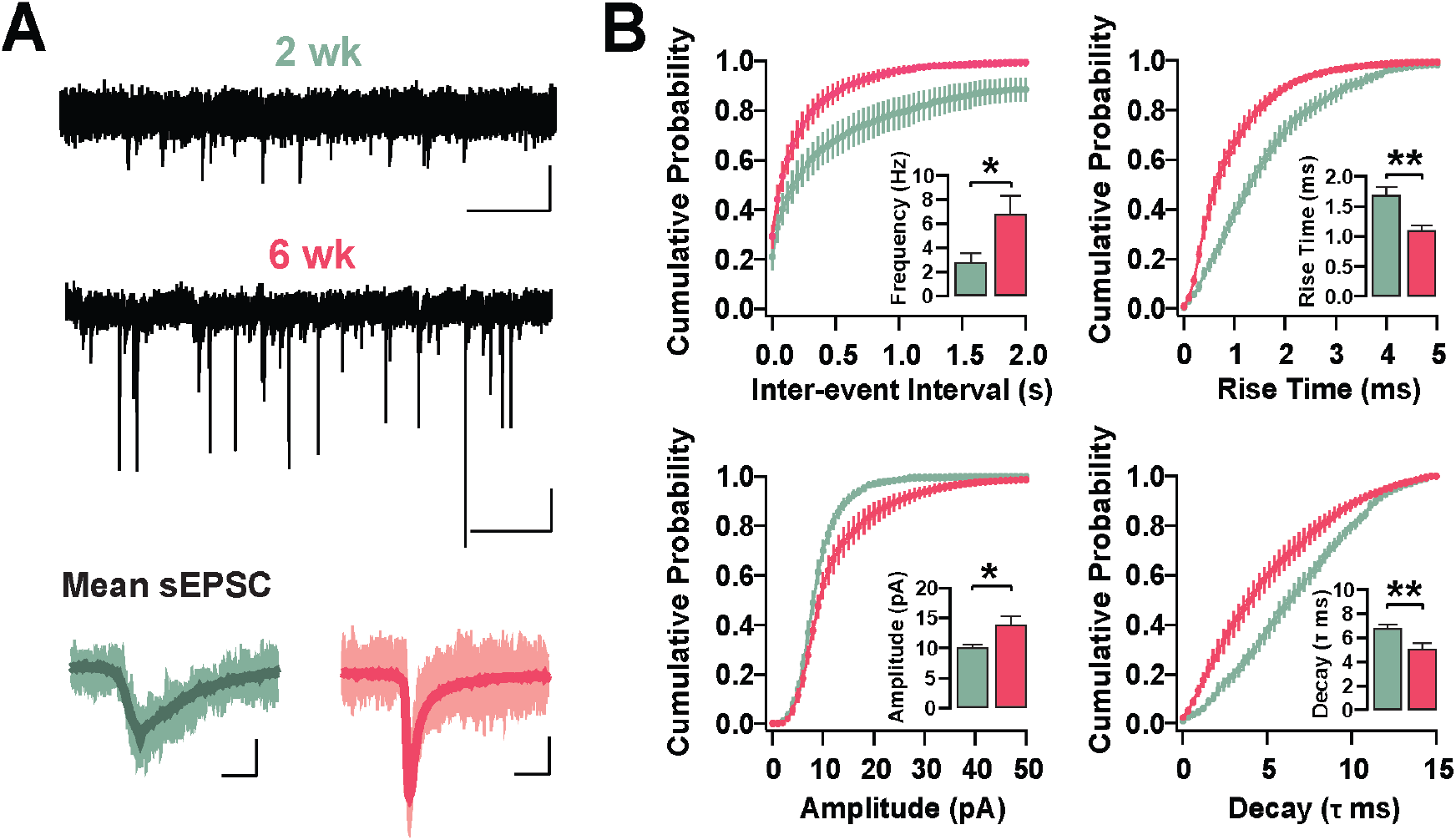
Changes in the kinetic properties of excitatory synapses in adult-born GCs **(A)** Top: representative voltage-clamp recordings at −70 mV showing spontaneous excitatory post-synaptic currents (sEPSCs) in GCs at 2 wk and 6 wk post-labeling (pink, 2 wk: n = 12 cells; green, 6 wk: n = 13 cells). The calibration is 10 pA and 500 ms. Bottom: average sEPSC waveforms from GCs at 2 and 6 wk, illustrating changes in amplitude, rise time, and decay with age. The calibration is 5 pA and 5 ms. **(B)** Average cumulative probability distributions for inter-event intervals, amplitude, rise time and decay for the sEPSCs in GCs at 2 and 6 wk post-labeling. Insets show the average frequencies and amplitudes (p = 0.02), and rise times and decays (p = 0.003) of sEPSCs at these two time points post-labeling.

**Supplementary Figure 2.**
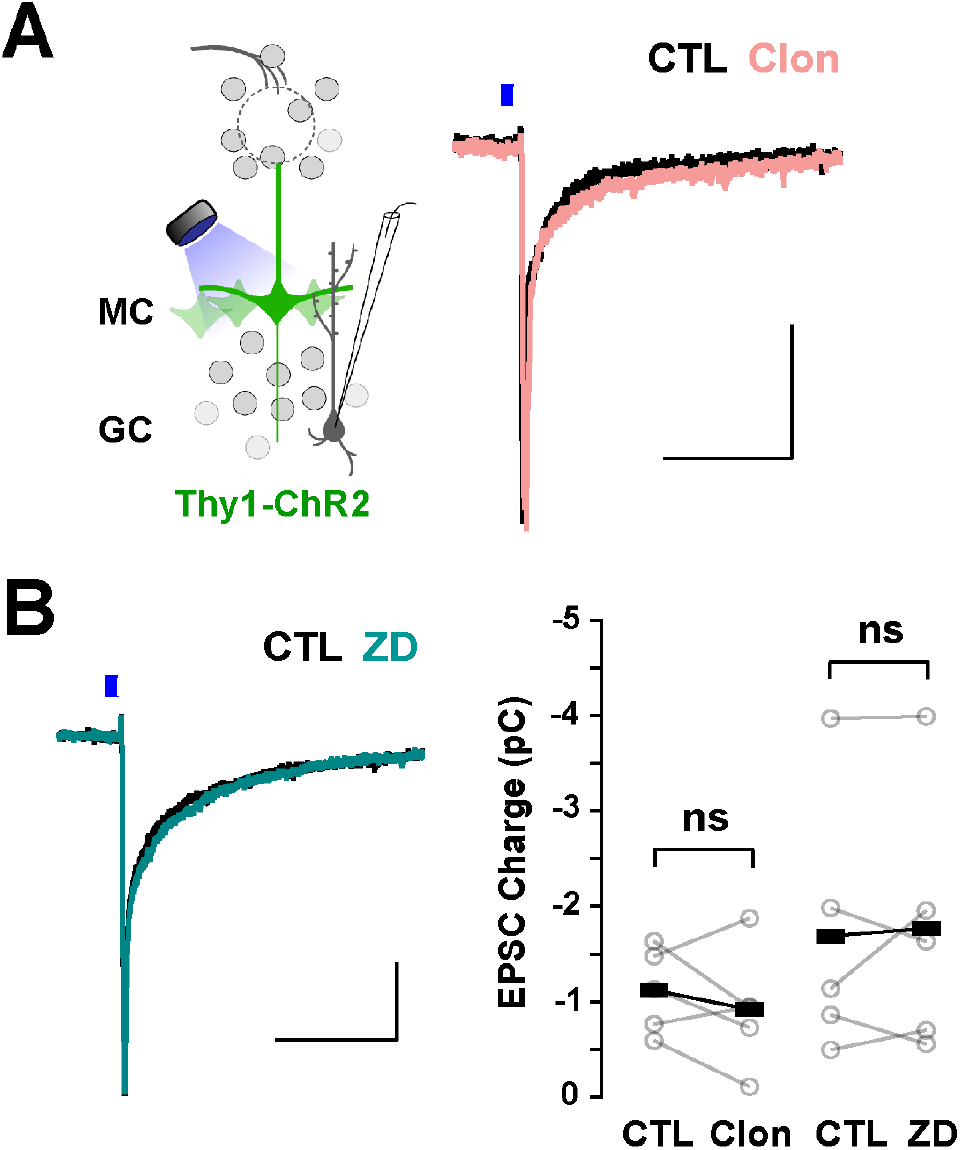
Blockade of I_h_ and activation of α2-ARs does not affect the MC to GC synapses **(A)** Left, diagram of the strategy used to elicit lateral inhibition. We recorded from GCs while optogenetically activating the MCs in OB slices from Thy1-ChR2 mice. Right, sample traces of light-elicited EPSCs in GCs at –70 mV during control (black) and in the presence of Clon (pink). The calibration bar is 10 pA and 200 ms. **(B)** Left, sample traces of light-elicited EPSCs in GCs during control (black) and in the presence of ZD. Right, summary plot of EPSC charge transfer showing no significant change in under both experimental conditions (Clon, n = 5 cells, p = 0.39; ZD, n = 5 cells, p = 0.73).

